# High-fidelity bioimage restoration via adversarial learning

**DOI:** 10.64898/2026.01.22.701051

**Authors:** Guillermo Rey-Paniagua, Dariusz Lachowski, Arrate Muñoz-Barrutia

## Abstract

Live-cell microscopy restoration is constrained by a trade-off between inference latency and texture preservation. While diffusion models provide high textural fidelity, the computational cost of iterative sampling currently limits their use in low-latency instrument feedback loops. Here, we present NAFNet GAN, a restoration framework that couples an activation-free backbone with a perceptual adversarial objective to enable high-throughput analysis. Unlike diffusion architectures, NAFNet GAN achieves an inference latency of **∼**110 ms for **1024 × 1024** inputs, potentially suitable for real-time instrument feedback loops. Across eight datasets ranging from STED nanoscopy to histopathology, the method achieves the lowest Learned Perceptual Image Patch Similarity (LPIPS) scores in 7 of 8 benchmarks while preserving structural coherence (e.g., MS-SSIM ***>* 0.968** in Cryo-EM), which facilitates reliable downstream analysis. Supported by performance benchmarks in the *AI4Life Denoising Challenge*, NAFNet GAN restores structural features from low-photon-budget acquisitions, maintaining the temporal resolution required for dynamic live-cell workflows.

## Introduction

Quantitative observation of biological systems requires balancing information retrieval against specimen integrity. While resolving subcellular architecture typically demands high signal-to-noise ratios (SNR), increasing photon flux drives phototoxicity and photobleaching, thereby imposing strict constraints on the observation of dynamic processes [1–3]. Beyond inherent shot noise, acquisitions are degraded by system-specific aberrations, including non-specific fluorophore binding, staining variability, and depth-of-field limitations. These degradations can hinder visual quality; they can affect downstream computational tasks, such as causing segmentation algorithms to fragment continuous membranes and tracking software to lose targets amidst stochastic variance [4, 5].

Consequently, as experimental designs increasingly necessitate low-photon budgets to preserve viability, restoration algorithms must recover quantitative signal from data dominated by Poisson–Gaussian noise and detector non-linearities [6–10]. Current restoration frameworks are often limited by a trade-off between computational efficiency and the preservation of high-frequency textures, which may impact their utility in high-throughput workflows [11, 12]. Regression-based networks (e.g., U-Nets, CARE) effectively minimize pixel-wise error but achieve this by averaging high-frequency spatial variance [13–15]. While statistically stable, this averaging smoothes high-frequency morphological details, such as cristae junctions or actin filaments, thereby limiting the precision of downstream quantitative phenotyping [11, 16].

Conversely, while generative diffusion models (DDPMs) provide high fidelity in textural recovery, their application to high-throughput workflows is challenged by inference latencies that complicate integration into instrument feedback loops, and the potential for stochastic sampling artifacts to introduce non-biological structural variations [12, 17, 18]. For quantitative biology, a restoration framework should prioritize morphological consistency without the computational overhead of iterative sampling. Here, we introduce NAFNet GAN, a morphology-preserving framework that couples a computationally efficient, activation-free backbone with a perceptual adversarial objective. Whereas regression models typically prioritize pixel-wise accuracy, often at the expense of texture, NAFNet GAN optimizes for deep feature similarity, aiming to reconstruct high-frequency spatial features relevant for morphological phenotyping [5, 19]. The architecture integrates a Nonlinear Activation-Free Network (NAFNet) generator [20], which maximizes signal propagation by replacing traditional gating non-linearities (e.g., Sigmoid or ReLU) with simple multiplication between feature-map splits. This architectural choice reduces computational complexity and preserves a more linear gradient flow, facilitating the recovery of subtle intensity variations and high-frequency textures characteristic of fluorescence microscopy. This design enables single-pass restoration (∼110 ms for 1024 × 1024 inputs), enabling direct integration into automated microscopy pipelines (Fig. 1).

**Fig. 1.**
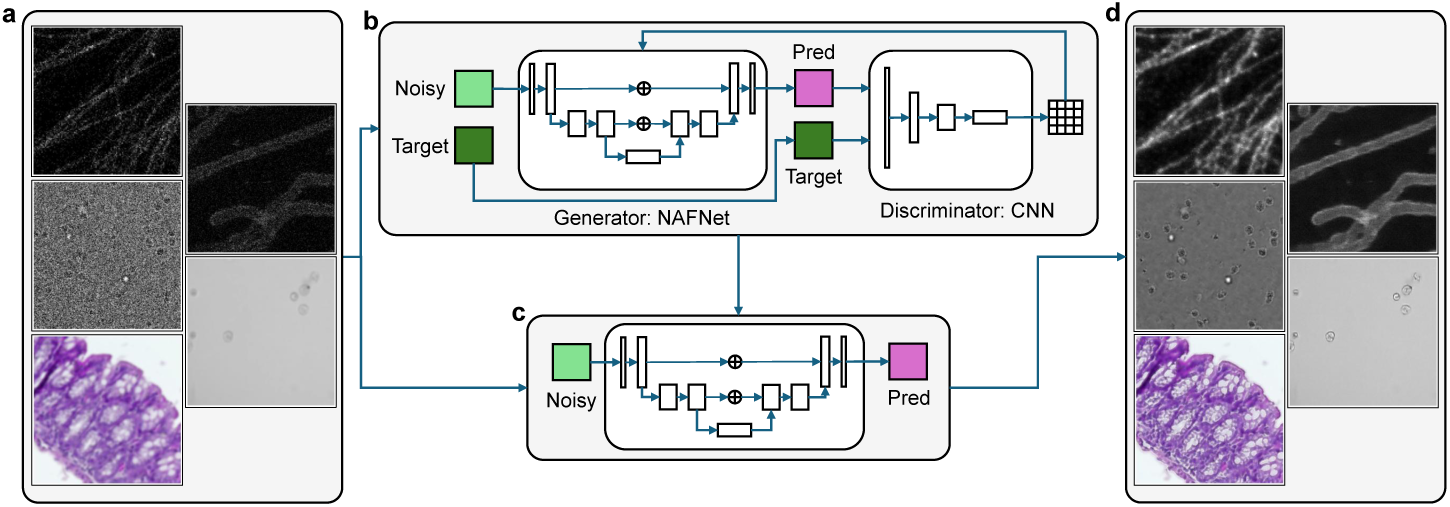
The NAFNet GAN framework. **a**, Representative samples illustrating low-SNR inputs across varying biological modalities. **b**, Schematic of the adversarial training pipeline. The Generator, built on a Nonlinear Activation-Free (NAFNet) backbone, predicts a restored image from the noisy input. The Discriminator classifies inputs between the restored prediction (Pred) and the ground truth (Target), enforcing textural consistency. The system is optimized via a composite loss combining pixel-wise fidelity and perceptual similarity. **c**, Inference workflow. Once trained, the Generator performs single-pass restoration on held-out data. **d**, Validation results demonstrating the recovery of cytoskeletal filaments and organelle morphology.

Our optimization strategy addresses the fundamental perception-distortion tradeoff by prioritizing *Learned Perceptual Image Patch Similarity* (LPIPS) over traditional pixel-wise metrics such as Mean Absolute Error (MAE). While MAE minimization forces the network to converge toward the conditional mean of the posterior distribution—typically resulting in an over-smoothed, blurry estimate—LPIPS penalizes discrepancies within the deep feature space [11, 21]. This allows NAFNet GAN to recover the high-frequency boundaries and structural textures that are conventionally attenuated by regression-based approaches.

We evaluated NAFNet GAN across eight datasets representing a broad range of signal-to-noise ratios, from Stimulated Emission Depletion (STED) nanoscopy to whole-tissue histopathology. Benchmarking indicates that our framework recovers high-frequency spatial features consistent with sub-pixel structures, where regression baselines tend to blur [14, 22–24] and diffusion models exhibit stochastic drift [12, 17, 25, 26]. These results suggest NAFNet GAN as a viable candidate for quantitative imaging workflows where both inference speed and morphological fidelity are essential.

## Results

### A Multimodal Validation Framework

The performance of NAFNet GAN was rigorously characterized using a multimodal validation framework encompassing eight datasets across seven imaging modalities (Fig. 2). To ensure an unbiased assessment of generalization, we partitioned each dataset into training and 25% held-out test subsets (Methods, Subsec. 1). The resulting validation suite samples a broad signal-to-noise spectrum, ranging from high-SNR sub-diffraction confocal nanoscopy (mean input MAE ≈ 0.035) to the photon-limited regime of Cryo-EM (mean input MAE ≈ 0.459; Extended Data Fig. 1a–d). For applications where experimental ground truth is inaccessible, we utilized unpaired datasets with parameterized noise profiles to simulate specific physical acquisition constraints (Fig. 2e–h and Methods, Subsec. 1).

**Fig. 2.**
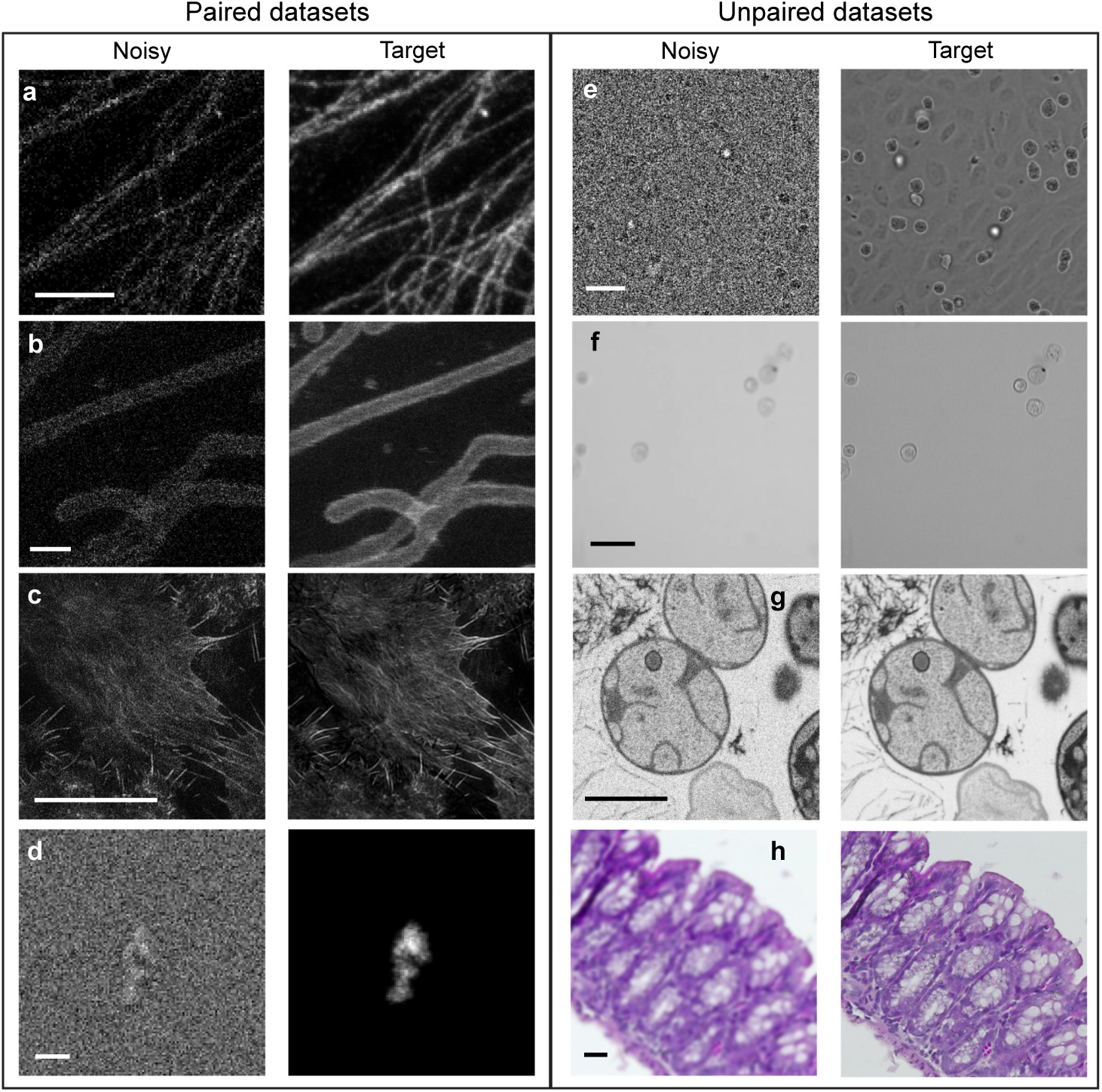
Cross-modal validation suite spanning diverse biological imaging modalities. The dataset is stratified into paired (supervised) and unpaired (synthetic degradation) regimes to test restoration across varying scales and noise profiles. **a**–**d**, Paired benchmarks containing experimentally acquired noisy inputs and high-SNR targets. **a**, Super-resolution STED microscopy of microtubules. **b**, STED microscopy of mitochondria. **c**, Confocal imaging of actin filaments. **d**, Cryo-EM of antibodies with simulated white Gaussian noise. **e**–**h**, Unpaired benchmarks where high-SNR acquisitions were degraded via synthetic noise/blur pipelines (Methods, Subsec. 2). **e**, Fluorescence microscopy of nuclei. **f**, Flow chamber imaging of cancer cells. **g**, FIB-SEM of *Gemmata obscuriglobus* bacteria. **h**, Brightfield histology of colonic tissue. **Scale bars: a**, **b**, 1 *µ*m; **c**, 10 *µ*m; **d**, 0.01 *µ*m; **e**, **f**, 50 *µ*m; **g**, 0.025 *µ*m; **h**, 1000 *µ*m.

Standardized nomenclature for these datasets follows the MODALITY-SAMPLE, DIMENSION, QUALITY-TARGET convention [27] (e.g., STED-C2DL-MTUB).

Technical specifications regarding acquisition parameters, noise modeling, and quantitative metrics—including Mean Absolute Error and MS-SSIM—are provided in the Methods (Subsec. 1) and Extended Data Fig. 1.

### Comparative baselines

We compared NAFNet GAN against representative architectures spanning regression, diffusion, and adversarial frameworks (Fig. 3). Pixel-wise regression baselines included CARE (U-Net) [13, 14] and Noise2Void [23]. To isolate the contribution of the adversarial loss from the architectural baseline, we also evaluated deep regression models: UNet-RCAN [19] and the NAFNet backbone [20] optimized exclusively for pixel-wise error. These were contrasted with generative approaches, specifically DDPM [26] (probabilistic diffusion) and Pix2Pix [28] (conditional GAN). All baselines were retrained on identical data partitions to ensure fair comparison (Methods, Subsec. 2).

**Fig. 3.**
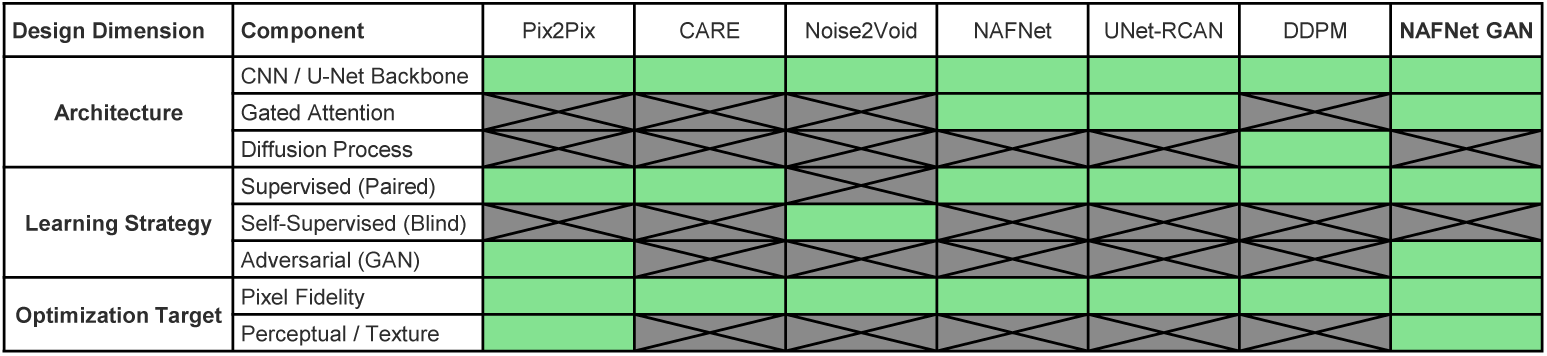
Functional taxonomy of bioimage restoration methods. Schematic classification of restoration frameworks based on backbone architecture, learning strategy, and optimization target. Methods are stratified into: (1) Regression models (CARE, Noise2Void) and Deep regression models (UNet-RCAN, standard NAFNet), which utilize pixel-wise loss functions; (2) Diffusion models (DDPM), characterized by iterative stochastic denoising; and (3) Adversarial frameworks (Pix2Pix, NAFNet GAN).

### Visual Recovery of High-Frequency Structures

NAFNet GAN balances pixel-level fidelity with perceptual similarity, restoring fine structures that are often smoothed by the averaging effects of regression models. Supplementing the reconstruction backbone with an adversarial objective (Methods, Eq. 1) mitigates the texture smoothing typical of standard regression networks. In mitochondrial nanoscopy (STED-C2DL-MITO), the model restores internal cristae topography with reduced structural residuals than baselines (Fig. 4, row 2), resulting in more distinct luminal boundaries that often appear blurred in MSE-optimized outputs. Similarly, in fluorescence microscopy (FLUO-N3DH-SIM), restoration sharpens nuclear envelopes, separating adjacent nuclei that appear spatially conflated due to focal blur in the raw input (Fig. 4, row 5).

**Fig. 4.**
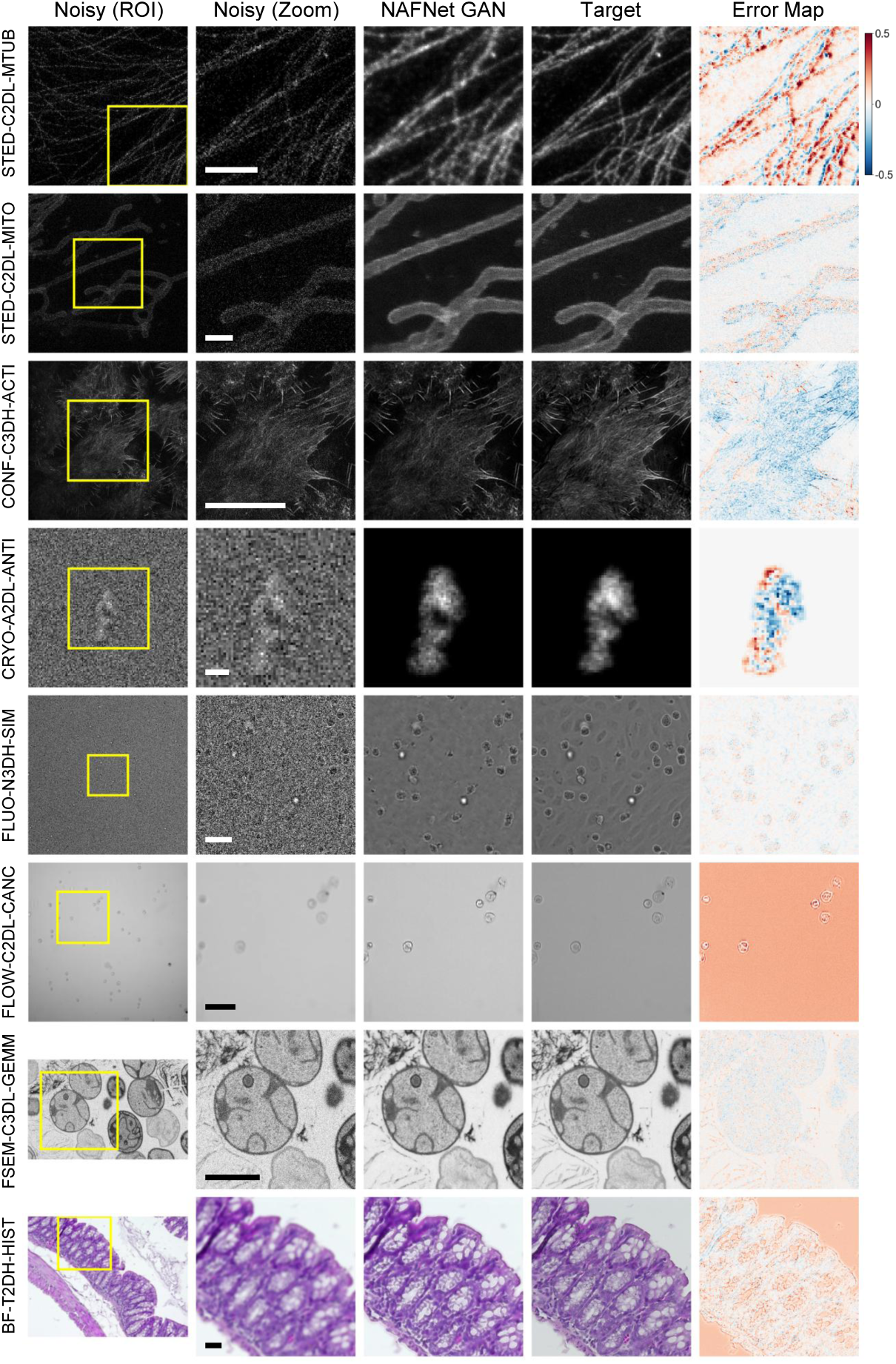
Visual assessment of restoration fidelity across biological scales. Representative validation results spanning diverse modalities (rows). Columns display (left to right): (1) Noisy (ROI): Full field-of-view input with the zoom region indicated in yellow. (2) Noisy (Zoom): Magnified view of the degraded region, highlighting specific artifacts (e.g., Poisson noise, blur). (3) NAFNet GAN: The model’s restoration, demonstrating recovery of high-frequency texture. (4) Target: High-SNR ground truth. (5) Error Map: Pixel-wise difference between the restoration and target. The map encodes zero error as white, with residuals diverging towards blue (Target *>* Restoration) or red (Restoration *>* Target). Scale bars (top to bottom): 1 *µ*m; 1 *µ*m; 10 *µ*m; 0.01 *µ*m; 50 *µ*m; 50 *µ*m; 0.025 *µ*m; 1000 *µ*m.

This fidelity extends to tissue-scale architectures. In histological sections (BF-T2DH-HIST), NAFNet GAN resolves crypt boundaries, aiding in the identification of morphological features to essential for more reliable histopathological assessment (Fig. 4, row 8). Analysis of residual error maps (Fig. 4, col. 5) shows that while adversarial optimization introduces a minor structural broadening in sub-diffraction features such as filaments (Fig. 4, row 1), the errors are predominantly stochastic and display minimal systematic geometric artifacts across diverse modalities.

### Comparative Assessment against State-of-the-Art Frameworks

Comparative benchmarking indicates three potential limitations in current methodologies (Fig. 5). First, deep regression-based models (NAFNet control, UNet-RCAN) tend to attenuate high-frequency biological signals. In Cryo-EM (CRYO-A2DL-ANTI), these architectures show reduced capability to distinguish globular antibodies from the noise floor, making target structures indistinguishable from background (Fig. 5, row 4). Similarly, in flow cytometry (FLOW-C2DL-CANC), baselines optimized for pixel-wise and adversarial fidelity (Noise2Void, Pix2Pix) exhibit limited capacity to recover distinct cellular boundaries (Fig. 5 row 6). Second, we observed artificial thickening of cytoskeletal targets (STED-C2DL-MTUB). Regression baselines produce filaments that are broadened compared to ground truth (Fig. 5, row 1); while this dilation minimizes pixel-wise error penalties (MAE) by optimizing spatial overlap, it constrains the spatial precision required for fine-grained filament segmentation. Finally, complex architectures such as DDPM and CARE exhibit spectral and spatial inconsistencies on the RGB Histology dataset (BF-T2DH-HIST), failing to fully compensate for the point-spread function and yielding residual blur and chromatic artifacts (Fig. 5, row 8).

**Fig. 5.**
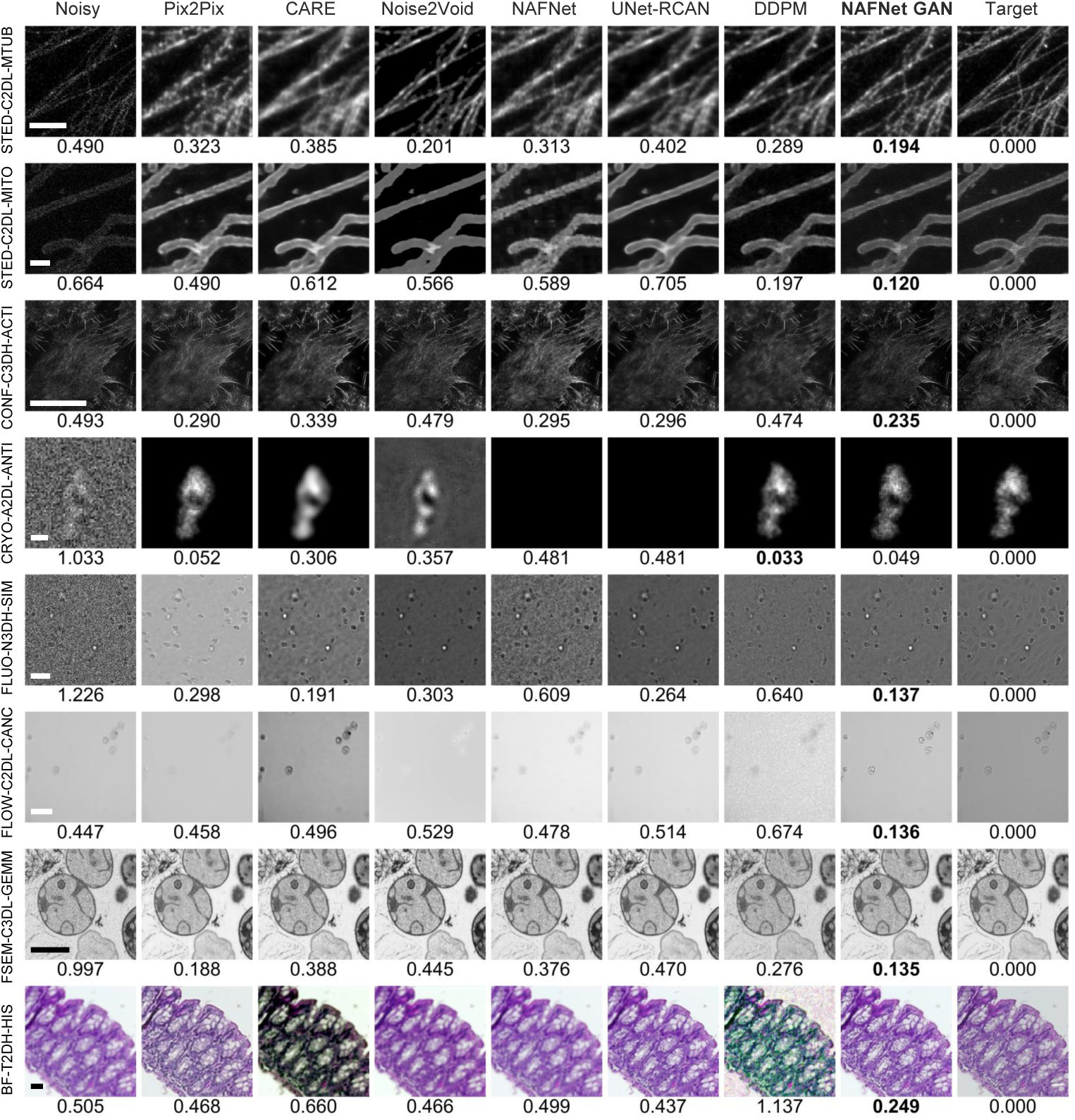
Qualitative comparison against established restoration methods. Restoration performance across eight diverse biological datasets (rows). Columns display outputs from representative frameworks. From left to right: (1) Noisy Input; (2) Pix2Pix (Adversarial baseline); (3–4) CARE, Noise2Void (Standard regression); (5–6) NAFNet, UNet-RCAN (Deep regression); (7) DDPM (Diffusion); (8) NAFNet GAN (Ours); (9) Target (Ground Truth). Values below each row indicate the LPIPS perceptual distance (lower is better) between the image above and the Target. Scale bars (top to bottom): 1 *µ*m; 1 *µ*m; 10 *µ*m; 0.01 *µ*m; 50 *µ*m; 50 *µ*m; 0.025 *µ*m; 1000 *µ*m.

### Statistical analysis of perceptual and photometric quality

Quantitative benchmarking illustrates the perception-distortion trade-off [11] (Fig. 6). While regression-based architectures (CARE, UNet-RCAN) minimize pixel-wise error (MAE, Eq. 6), they average high-frequency details, causing texture oversmoothing and elevated perceptual error (LPIPS, Eq. 3).

**Fig. 6.**
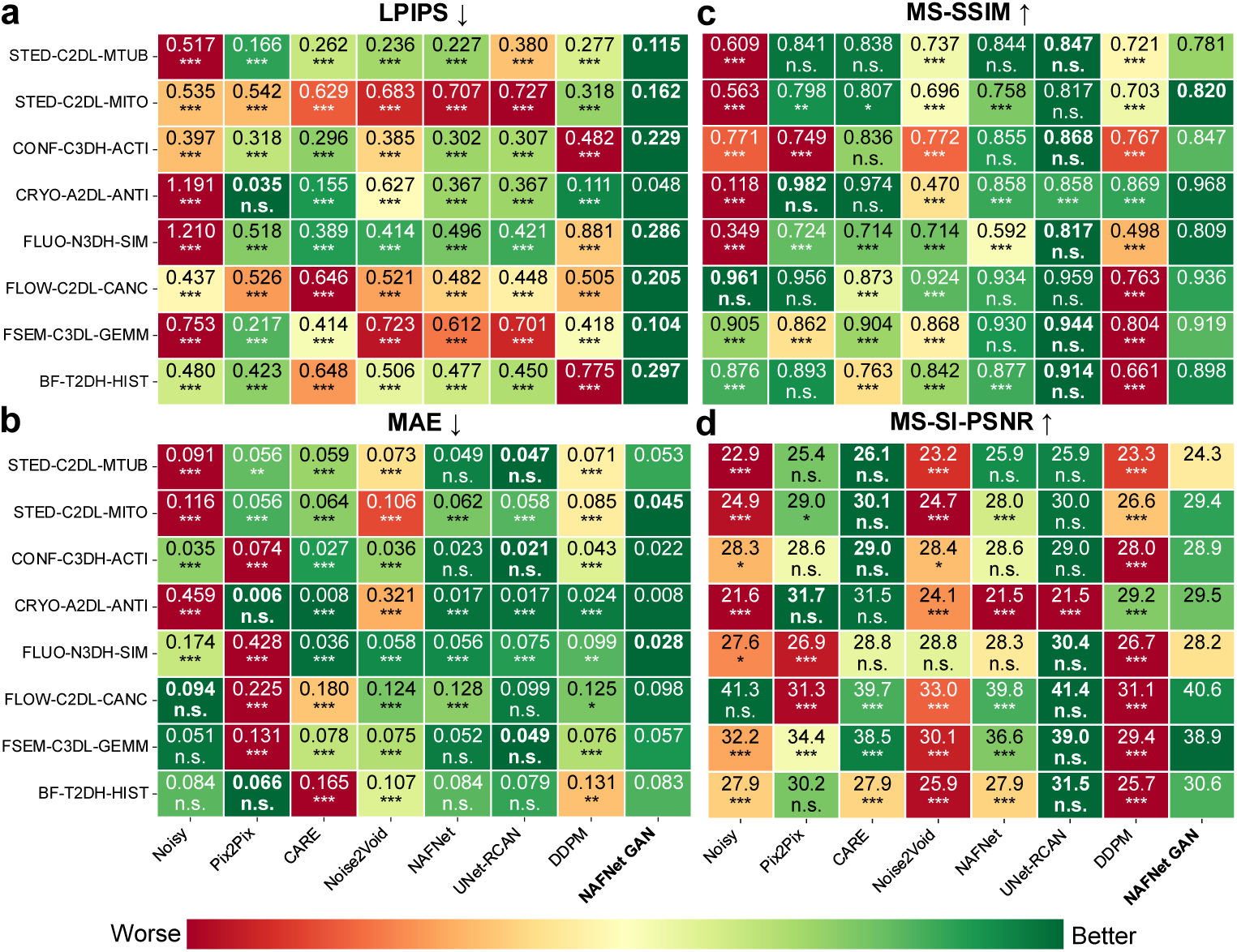
Quantitative benchmarking of restoration performance. Heatmap summary comparing NAFNet GAN against six baseline architectures across eight datasets. Columns represent evaluation metrics defined in Methods (Subsec. 2): **a**, Learned Perceptual Image Patch Similarity (LPIPS), quantifying textural fidelity (lower is better); **b**, Mean Absolute Error (MAE), measuring pixel-wise precision (lower is better); **c**, Multi-Scale Structural Similarity Index (MS-SSIM) (higher is better); and **d**, Multi-Scale Signal-Invariant PSNR (MS-SI-PSNR) (higher is better). Color scale indicates relative ranking (green: optimal; red: suboptimal). Statistical significance against the second-best method was determined by one-sided Welch’s t-test (* *P <* 0.05; ** *P <* 0.01; *** *P <* 0.001; n.s., not significant).

NAFNet GAN addresses the perception-distortion trade-off. Statistical analysis via one-sided Welch’s *t*-test demonstrates that our framework minimizes perceptual distance (*P <* 0.001) in 7 of 8 datasets compared to top-performing baselines (Fig. 6). In histopathology (BF-T2DH-HIST), for instance, NAFNet GAN reduced perceptual error by 30% (LPIPS 0.297 vs. 0.423; *P <* 0.001), recovering crypt boundaries (MAE 0.066) obscured by regression models. This observation supports the view that minimizing perceptual distance serves as a more robust proxy for structural recovery than absolute pixel-wise fidelity.

The prioritization of perceptual metrics may introduce deviations from absolute pixel-wise accuracy, as observed in STED-C2DL-MTUB, where NAFNet GAN exhibits a higher MAE than UNet-RCAN (Fig. 6b). Residual analysis (Fig. 4) suggests that these photometric deviations are largely confined to non-structural background regions. While regression losses converge to the conditional mean, leading to boundary blur, the adversarial objective (Eq. 1) preserves the high-frequency texture required to recover adjacent structures. NAFNet GAN maintains high structural fidelity and signal invariance under linear intensity shifts, as evidenced by a mean MS-SI-PSNR (Eq. 5) of 29.4 on the STED-C2DL-MITO dataset. This stability supports the preservation of local gradients necessary for downstream segmentation.

### Efficiency and Throughput

High-throughput microscopy requires balancing reconstruction quality with inference latency across variable field-of-views (FOVs). We benchmarked inference speeds across dimensions ranging from 128 × 128 px (Cryo-EM) to 1360 × 1024 px (Histology) (Extended Data Fig. 2).

Lightweight regression architectures (CARE, Noise2Void) consistently yield high throughput, operating in the sub-100 ms regime even for megapixel-scale inputs (Extended Data Fig. 2). However, their utility is limited by the loss of high-frequency texture. Conversely, increasing regression depth yields marginal performance gains; UNet-RCAN requires ∼740 ms for a 1024 × 1024 frame (CONF-C3DH-ACTI), yet retains smoothing artifacts.

Probabilistic diffusion models (DDPMs) exhibit significant latency scaling. While effective at texture recovery, their iterative sampling increases quadratically with resolution. In high-resolution regimes (e.g., 1024 × 1024 px), DDPM requires ∼3.5 s per frame—approximately 30× slower than NAFNet GAN—potentially precluding its application for dynamic acquisition. NAFNet GAN mitigates this bottleneck through single-pass generator inference. It demonstrates linear scalability, processing small FOVs (e.g., 128 × 128 px Cryo-EM) in ∼5 ms and large FOVs (1024 × 1024 px) in ∼113 ms. This profile enables high-content screening with perceptual fidelity competitive with diffusion models.

### Robustness Across Degradation Regimes

Beyond computational efficiency, reliability under varying signal qualities is a critical requirement for deployment. We defined the operational limits of NAFNet GAN via robustness analysis across varying degradation levels (Methods, Subsec. 2), assessing fidelity from ideal inputs to regimes of severe corruption (*σ* = 0.10 noise; *σ*_blur_ = 5 blur; Extended Figs. 3–6).

The model separates signal from noise while preserving underlying morphology in high-SNR regimes. In the “No Noise” control condition (Extended Figs. 3–4, top row), NAFNet GAN maintains high input fidelity (*LPIPS* ≈ 0.000). This indicates that the model prioritizes signal preservation, substantially reducing the risk of stochastic artifacts commonly observed in generative inference on clean data.

Under severe degradation, NAFNet GAN maintains perceptual consistency (LPIPS *<* 0.5) even at the upper bounds of the training envelope. This stability was found to be statistically significant (*P <* 0.001; Extended Figs. 5, 6); our framework outperformed regression (CARE, Noise2Void) and deep regression (NAFNet, UNet-RCAN) baselines across most tested noise and blur intensities. Whereas regression models displayed characteristic over-smoothing and diffusion models (DDPM) exhibited instability under severe noise conditions, NAFNet GAN preserved structural coherence, outperforming the adversarial baseline (Pix2Pix). This robustness was further supported by performance on the blind test set of the *AI4Life Microscopy Denoising Challenge*, where our method achieved the highest performance benchmarks in three of four categories (Extended Data Fig. 7), demonstrating generalizability beyond internal benchmarks.

## Discussion

Computational microscopy aims to maximize information retrieval from limited photon flux, enabling the observation of sensitive biological processes constrained by phototoxicity [1, 2]. However, many existing restoration methods are characterized by a tension between the statistical stability of regression models—which maximize signal-to-noise ratios by averaging out high-frequency texture [11, 14]—and the perceptual resolution of diffusion models, which impose substantial computational costs and introduce stochastic variability [25, 29].

We propose addressing this limitation through architectural optimization. By coupling a computationally efficient, activation-free backbone with a morphology-consistent adversarial objective, NAFNet GAN seeks to balance these competing constraints. We demonstrate the restoration of features consistent with sub-diffraction geometry at latencies compatible with live imaging. This approach aims to address the trade-off between quantitative fidelity and perceptual realism, while reducing the inference artifacts often associated with unconstrained generative approaches.

### Morphological Fidelity versus Photometric Precision

The adoption of generative models in quantitative workflows has been limited by the potential for unfaithful structural reconstruction [18]. Our analysis suggests that current caution regarding generative models may stem from a reliance on pixel-wise metrics (e.g., PSNR) that prioritize photometric conservation over structural recoverability.

In low-SNR regimes, such as Cryo-EM (CRYO-A2DL-ANTI), regression-based optimization minimizes global error by converging to the conditional mean. While mathematically stable, this smooths high-frequency particle signals, limiting data utility for fine-grained structural analysis. This was demonstrated in the outputs of the baseline regression models, where such features were effectively suppressed as background noise. In contrast, NAFNet GAN utilizes the discriminator as a learned textural prior, promoting the reconstruction of features consistent with the target distribution even when the signal is obscured by noise.

While this approach emphasizes local contrast over absolute photometric fidelity—resulting in minor intensity offsets under severe Gaussian blurring (Extended Figs. 3, 4)—it preserves the morphological features (boundaries, gradients) necessary for characterization (Eq. 1). For downstream tasks, such as segmentation or particle counting, this gradient recovery improves accuracy, whereas excessive smoothing obscures morphological cues, as evidenced in Fig. 5 row 6.

### Cross-Modal Generalization

A significant challenge for deep learning adoption in core facilities can be limited generalization—the tendency of algorithms trained on specific structures to fail on distinct morphologies. The consistent performance across eight imaging modalities suggests that a unified, activation-free architecture can generalize across diverse contrast mechanisms without the need for dataset-specific architectural tuning.

The consistent performance across diverse modalities suggests that the discriminator identifies cross-modal structural priors, such as filament continuity (see Fig. 4, row 1), rather than overfitting to specific noise signatures. This robustness, supported by the superior performance benchmarks in the *AI4Life Denoising Challenge* (Extended Data Fig. 7), supports the feasibility of a unified restoration pipeline, reducing the operational burden of custom-trained networks.

This generalization furthermore extends to spectral integrity. Whereas conventional architectures often process multi-channel inputs independently, leading to spectral decoupling associated chromatic fringing (Fig. 5, bottom row). Conversely, NAFNet GAN processes the multi-channel volume as a coherent entity. By enforcing consistency across the spectral dimension, the model maintains the chromatic balance required for diagnostic tasks without requiring channel-specific retraining [30, 31].

### Signal Preservation in Ideal Acquisition Regimes

Successful integration of generative models into quantitative workflows requires demonstrable signal preservation. As demonstrated in our robustness analysis (Extended Figs. 3, 4), NAFNet GAN exhibits conservative behavior in high-signal regimes. When presented with clean inputs, the model maintains high input fidelity, evidenced by LPIPS values approaching zero in “No Noise” conditions.

This property minimizes the potential for artifact introduction into valid biological signals, particularly in applications lacking paired ground truth. In contrast to diffusion models, which iteratively reconstruct the input regardless of quality, our architecture leverages global residual learning. By constraining the network to predict only the noise component, the backbone maintains a near-zero residual state in the absence of degradation. This ensures that NAFNet GAN functions as a conditional filter, suitable for variable imaging conditions without compromising high-quality frames.

### Implications for Real-Time Adaptive Microscopy

The computational efficiency of NAFNet GAN facilitates the transition from offline post-processing to closed-loop feedback. While diffusion models offer high-fidelity texture recovery, their iterative sampling mechanism limits their utility in dynamic acquisition loops.

With a mean inference latency of ∼113 ms for 1024 × 1024 inputs, NAFNet GAN enables adaptive microscopy workflows. This latency profile could potentially support adaptive microscopy workflows where real-time restoration of preview streams assists in the automated identification of transient biological events—such as mitosis onset—and trigger high-resolution capture only when necessary. By improving restoration quality at high speeds, this approach shifts a portion of the experimental burden from physical photon acquisition to computational processing.

By reducing inference latency while maintaining structural fidelity, NAFNet GAN provides a feasible pathway for extending live-cell imaging durations via low-dose acquisition protocols. We demonstrate that the recovery of features consistent with sub-diffraction structures is achievable with a reduced risk of stochastic artifacts compared to standard regressive and adversarial architectures, and with significantly lower computational overhead than iterative diffusion-based models. These efficiency gains may help transition image restoration from an offline post-processing task toward a low-latency feedback component, contributing to the development of autonomous, photon-efficient microscopy workflows designed to minimize phototoxicity while preserving data utility.

## Methods

### Dataset preparation and benchmarking

To evaluate generalizability across biological scales, we compiled a diverse suite of eight datasets spanning seven imaging modalities (Fig. 2), stratified into paired (experimentally acquired ground truth) and unpaired (synthetically degraded) regimes. Detailed acquisition parameters and statistics are provided in Extended Data Fig. 1.

To ensure unbiased evaluation, data from each dataset was partitioned into training (60%), validation (15%), and testing (25%) subsets. This protocol prevents data leakage, ensuring that no spatial patches from the same biological sample were shared between optimization and evaluation phases.

#### Paired datasets

Morphological reconstruction was assessed using two STED datasets [26]: STED-C2DL-MTUB (1,319 images, 256 × 256 px) containing dense microtubules degraded by Poisson–Gaussian noise, and STED-C2DL-MITO (345 images, 600×600 px) capturing mitochondrial cristae. Confocal restoration was evaluated using CONF-C3DH-ACTI [32] (774 images, 1024 × 1024 px), characterizing low-noise filamentous recovery. Low-SNR performance was benchmarked on CRYO-A2DL-ANTI [33], comprising 100,000 synthetic Cryo-EM images (128 × 128 px) with severe additive white Gaussian noise (mean MS-SSIM ∼0.118).

#### Unpaired and microscopic datasets

For regimes lacking experimental ground truth, we utilized FLUO-N3DL-SIM [34] (77 images, 1024 × 1024 px) to test deconvolution of focal blur, and FLOW-C2DL-CANC [35] (307 images, 1024 × 1024 px) to assess motion blur recovery in flow cytometry.

High-fidelity texture recovery was evaluated on FSEM-C3DL-GEMM [36] (470 FIB-SEM crops, 1600×1000 px) and BF-T2DH-HIST [37] (50 H&E histology scans, 1360× 1024 px), the latter serving as the sole RGB benchmark for chromatic fidelity.

### Dynamic degradation strategy

For unpaired datasets, we implemented an online augmentation pipeline. During training, inputs were dynamically degraded using one of three regimes selected with equal probability: (1) identity mapping; (2) additive Gaussian noise sampled uniformly from *σ* ∈ [0.01, 0.1]; or (3) Gaussian blur with kernel widths *σ_blur_* ∈ [2, 5]. This stochastic degradation pipeline models a range of heterogeneous noise floors, ensuring the network learns robust restoration features rather than over-fitting to specific acquisition noise priors [10].

### Network Architecture

#### Generator (NAFNet backbone)

The framework implements a conditional Generative Adversarial Network (cGAN). The generator incorporates a Nonlinear Activation-Free Network (NAFNet) backbone [20], configured as a four-level encoder–decoder with a base channel width of 16. This four-level configuration was chosen to maintain a balance between feature extraction capacity and inference latency, as supported by architectural ablation studies (Extended Data Fig. 8). The architecture replaces nonlinear activation functions with a Simple Gate mechanism (*Y* = *X*1 ⊙ *X*2) to preserve information flow through the network layers. Structural stability is supported via Simplified Channel Attention (SCA) [38] and global residual learning [40], constraining the model to approximate the residual noise component. Upsampling is performed via PixelShuffle [39].

#### Discriminator

The discriminator employs a PatchGAN architecture with a five-stage downsampling pathway. This design provides a receptive field tailored to sub-cellular features, facilitating the penalization of local structural discrepancies during the adversarial training phase.

### Loss functions

#### Primary objective

The NAFNet GAN model minimizes a composite objective function:

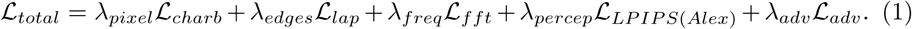

where L*_charb_* denotes the Charbonnier pixel-wise loss for robust pixel alignment, and L*_lap_* and L*_fft_* enforce consistency in the spatial gradient and spectral domains, respectively. The perceptual objective, L*_LP IP S_*_(*Alex*)_, utilizes a pre-trained AlexNet backbone to evaluate deep-feature similarity. While originally trained on natural images, this metric serves here as a high-dimensional proxy for textural complexity, capturing feature correlations often overlooked by traditional pixel-wise metrics [21]. The weighting coefficients (*λ_pixel_* = 0.5*, λ_edges_* = 2*, λ_freq_* = 0.5*, λ_percep_* = 3*, λ_adv_* = 0.5) were determined empirically via grid search on the validation set to reconcile the requirements for global structural alignment and fine-grained texture reconstruction.

#### Challenge baseline

For the *AI4Life Denoising Challenge 2025*, we adapted the objective function to maximize Structural Integrity PSNR (SI-PSNR), the competition’s primary metric.

This formulation emphasizes structural and perceptual alignment by incorporating a VGG-19 backbone (*λ_percep_* = 100)[21] and a high-weight Structural Similarity term (*λ_SSIM_* = 10):

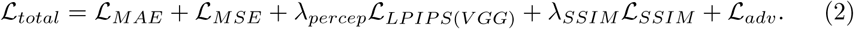

We combined these terms with standard pixel-wise error (L*_MAE_,* L*_MSE_*) and adversarial regularization to ensure robust performance across the challenge benchmarks.

### Training protocol

Training was conducted on a workstation equipped with a single NVIDIA GeForce RTX 5060 Ti (16 GB vRAM) and an AMD Ryzen 7 5800X CPU, using Python as the programming language. Input volumes were dynamically normalized to [0, 1]. During training, we applied the stochastic degradation strategy detailed in Methods Subsec. 2, alongside random batch cropping to 50% of the original spatial dimensions. For fair validation, we employed a deterministic degradation protocol comprising 40% Gaussian noise (*σ* = 0.05), 40% Gaussian blur (*σ_blur_* = 3), and 20% uncorrupted images. We trained an independent model for each of the eight datasets to ensure modality-specific optimization.

Optimization utilized AdamW (*β*_1_ = 0.5, *β*_2_ = 0.999, weight decay 1 × 10*^−^*^2^) with an initial learning rate of 1 × 10*^−^*^3^ and a cosine annealing schedule applied during the final 20% of iterations. Training duration varied by dataset size: 50 epochs (CONF-C3DH-ACTI), 30 epochs (CRYO-A2DL-ANTI), 300 epochs (BF-T2DH-HIST), and 100 epochs for remaining datasets. Reference baselines (Pix2Pix, CARE, Noise2Void, NAFNet, UNet-RCAN, DDPM) were retrained using their official implementations.

To obtain inference times, the GPU was constrained to 4 GB vRAM for a fair comparison (see Extended Data Fig. 2).

### Quantitative evaluation metrics

Metrics were computed on data normalized to [0, 1] in Python.

#### Learned Perceptual Image Patch Similarity (LPIPS)

Computed using the official AlexNet backbone [21], LPIPS calculates the weighted Euclidean distance between deep feature activations:

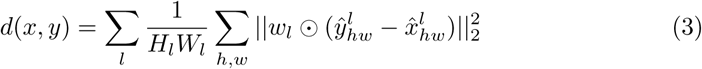

#### Multi-Scale Structural Similarity Index (MS-SSIM)

Calculated over a three-scale pyramid (*M* = 3) with a window size of 7 pixels and weights *β_j_*= *γ_j_* = {0.25, 0.50, 0.25} [41]:

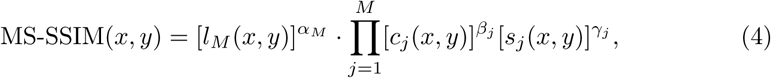

#### Multi-Scale Signal-Invariant PSNR (MS-SI-PSNR)

Computed across four spatial scales (*k* = 1 … 4) with weights *ω_k_* = {0.0448, 0.2856, 0.3001, 0.3695}. At each scale, the restored image is linearly aligned to the ground truth via least-squares regression to minimize global photometric error before PSNR calculation [42]:

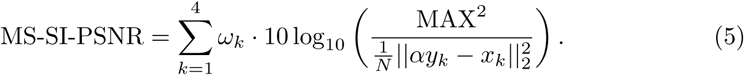

#### Mean Absolute Error (MAE)

Calculated as the pixel-wise *L*_1_ distance between the restoration and ground truth:

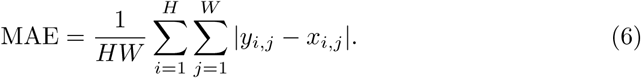

### Statistical analysis

To assess the significance of performance improvements—specifically comparing the NAFNet GAN baseline against the state-of-the-art, ablation variants, and the optimization strategies—we employed a one-sided Welch’s t-test. This test was selected to account for unequal variances in error distributions across the heterogeneous imaging datasets. Statistical significance was defined as *P <* 0.05 [43]. All statistical computations were performed using the scipy.stats library in Python.

## Declaration of Generative AI usage

Large language models (Gemini, Google) were used solely for copy-editing and syntax refinement. All scientific content and data analysis were verified by the authors.

## Data availability

The datasets generated and/or analyzed during the current study are available in the Zenodo and EMPIAR repositories: STED-C2DL-MTUB/MITO [44, 45], CONF-C3DH-ACTI [32], CRYO-A2DL-ANTI [46], FLUO-N3DL-SIM [34], FLOW-C2DL-CANC [47], FSEM-C3DL-GEMM [36], and BF-T2DH-HIST [37].

## Code availability

Source code is available on GitHub (https://github.com/GolpedeRemo37/ NAFNet-GAN-Perceptual-and-Real-time-Restoration-of-Biological-Structure).

## Acknowledgements

This work was partially supported by the Ministerio de Ciencia, Innovación y Universidades, Agencia Estatal de Investigación, MCIN/AEI/10.13039/501100011033/, under grants PID2023-152631OB-I00 and AIA2025-164165-C41, co-financed by European Regional Development Fund (ERDF), “A way of making Europe”. Additionally, Dariusz Lachowski was supported by Consejeŕıa de Educación, Ciencia y Universidades, Comunidad de Madrid Grant 2023-T1/SAL-GL-29109. We thank the AI4Life Microscopy Challenge organizers for hosting and evaluating the submissions.

## Author contributions

Conceptualization: all authors. Experiments: G.R-P. Writing—initial outline: all authors. Writing—original draft: all authors. Writing—review and editing: all authors. Illustrations: G.R-P. Supervision: D.L. and A.M-B. Funding acquisition: D.L. and A.M-B.

## Extended Data

**Extended Data Fig. 1.**
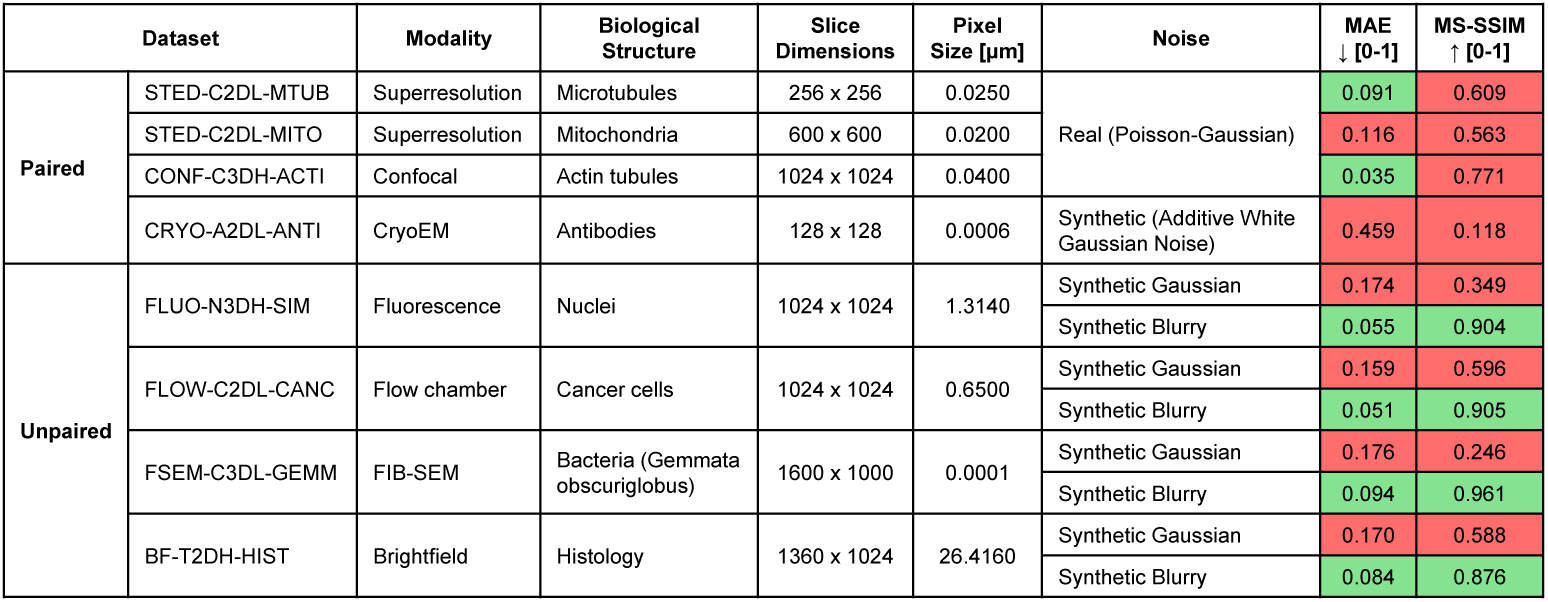
Dataset characteristics and difficulty stratification. Summary statistics for the eight validation datasets. Columns display: pairing status, standardized nomenclature, microscopy modality, biological target, image dimensions (*H*), pixel size (*µ*m), noise characteristics, and mean input metrics (MAE and MS-SSIM). Color scales indicate relative restoration difficulty (green: easier/higher quality; red: harder/lower quality), with thresholds set at 0.1 for MAE and 0.8 for MS-SSIM.

**Extended Data Fig. 2.**
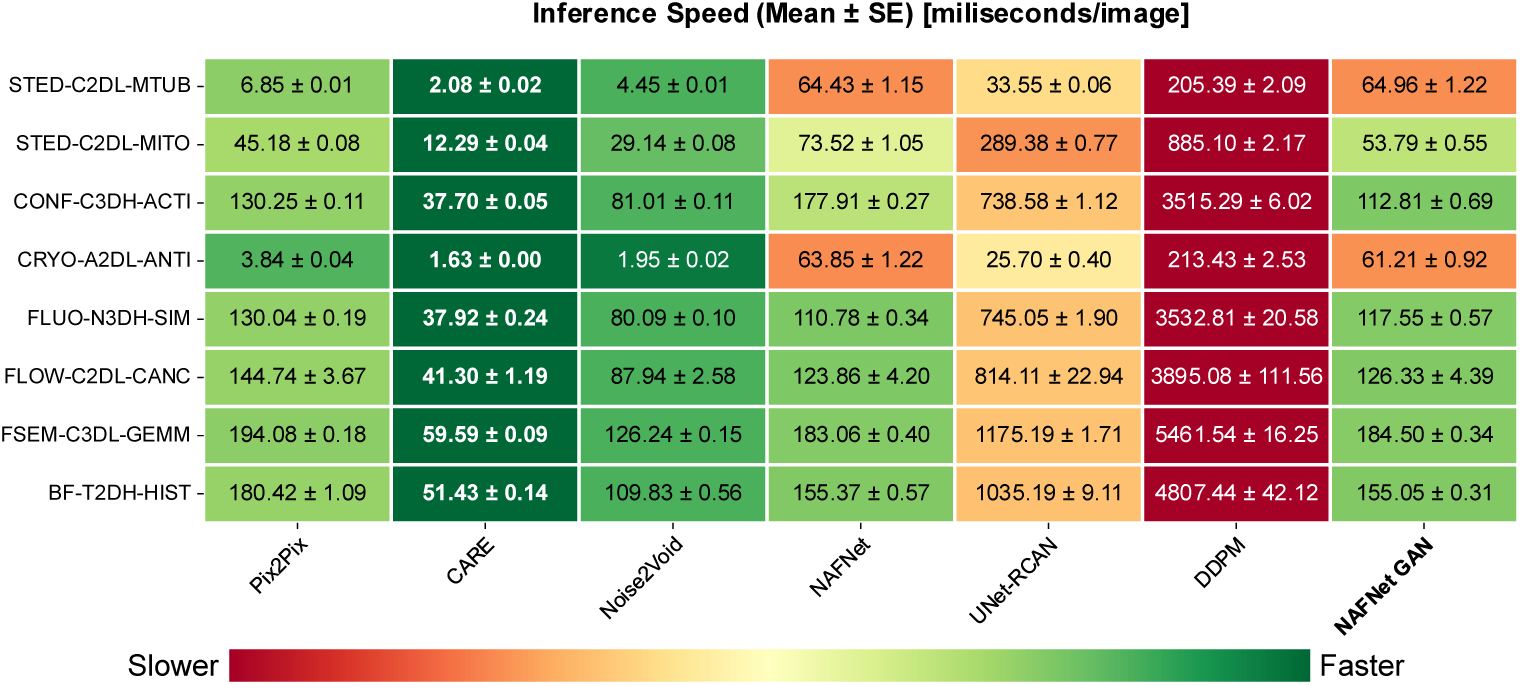
Inference latency benchmarking across variable acquisition dimensions. Comparative analysis of inference speed (miliseconds per image) versus input dimension. Architectures were evaluated on image sizes ranging from 128 × 128 px (Cryo-EM) to 1360 × 1024 px (Histology) under a constrained computational budget (4 GB VRAM). NAFNet GAN demonstrates efficient scalability, maintaining milisecond-scale inference (∼112.81 ms for 1024 × 1024 px) where diffusion models (DDPM: ∼3515.29 ms) become computationally prohibitive. Values represent mean ± s.e.m. (*n* = 200 images, or all available images for smaller datasets).

**Extended Data Fig. 3.**
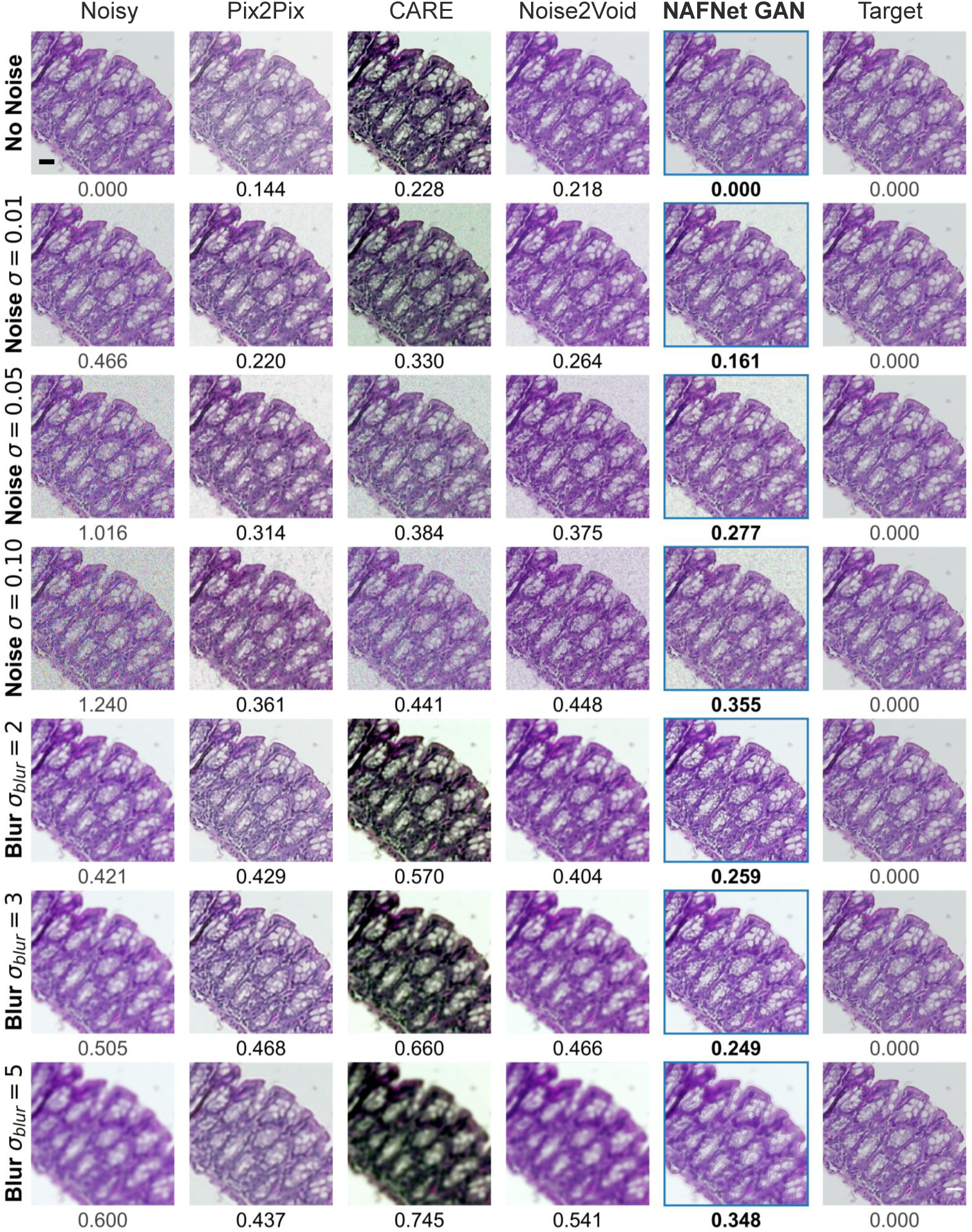
Restoration stability under varying degradation severities (Standard Baselines). Visual comparison of NAFNet GAN against adversarial (Pix2Pix) and standard regression (CARE, Noise2Void) baselines on the BF-T2DH-HIST dataset. Rows correspond to increasing degradation severities: ”No Noise” (Control), increasing Gaussian noise (*σ* ∈ {0.01, 0.05, 0.10}), and increasing Gaussian blur (*σ*_blur_ ∈ {2, 3, 5}). Values indicate LPIPS perceptual distance (lower is better). Note the strict signal preservation in the ”No Noise” condition (Row 1), confirming the model does not hallucinate structures on high-fidelity inputs. **Scale bar:** 1000 *µ*m.

**Extended Data Fig. 4.**
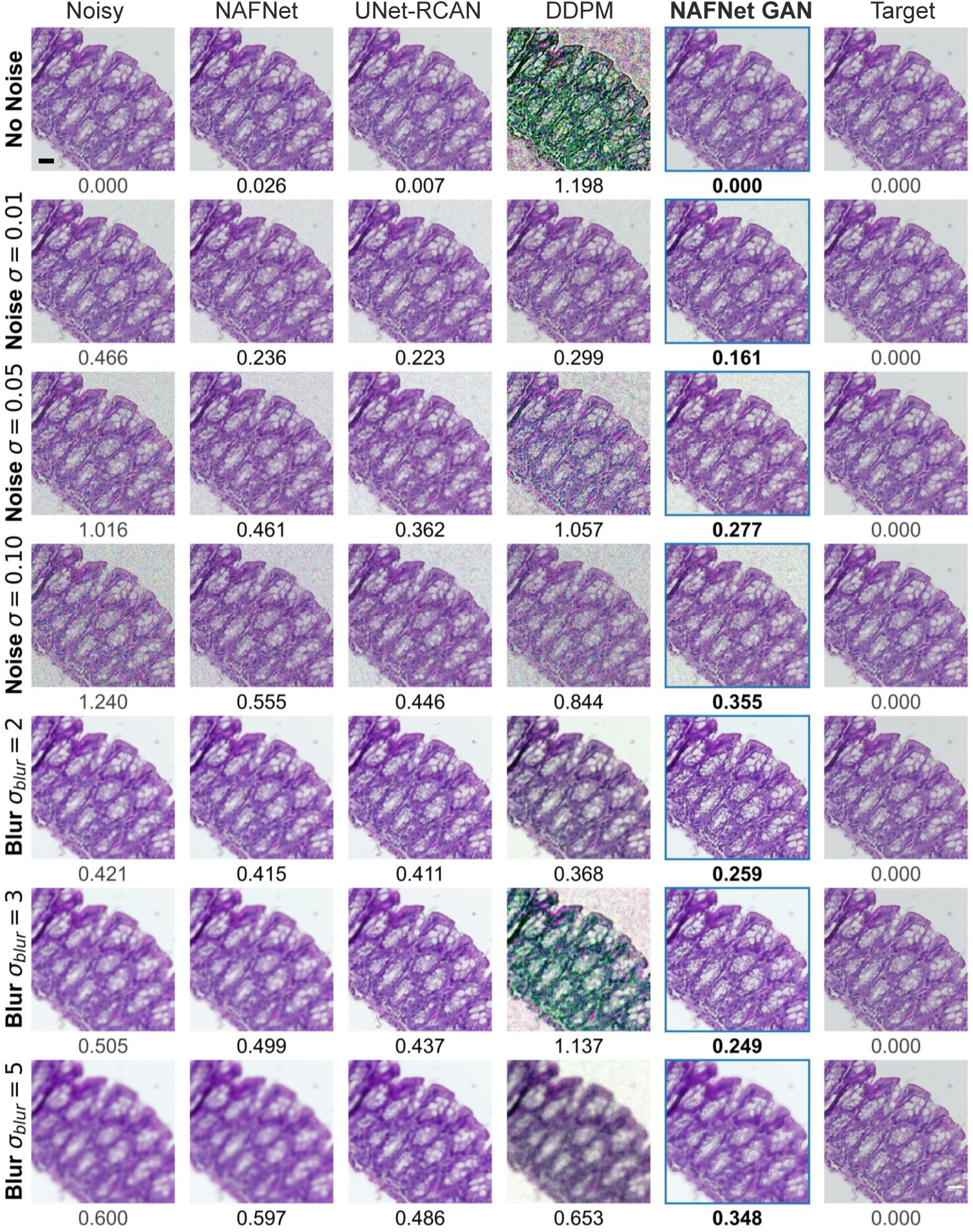
Restoration stability under varying degradation severities (Advanced Models). Continuation of the stability analysis on BF-T2DH-HIST, comparing NAFNet GAN against deep regression (NAFNet, UNet-RCAN) and diffusion (DDPM) architectures. Degradation conditions match Extended Data Fig. 3. NAFNet GAN maintains structural consistency without the texture collapse of regression models or the high-frequency artifacts observed in diffusion models under severe noise. Values indicate LPIPS score. **Scale bar:** 1000 *µ*m.

**Extended Data Fig. 5.**
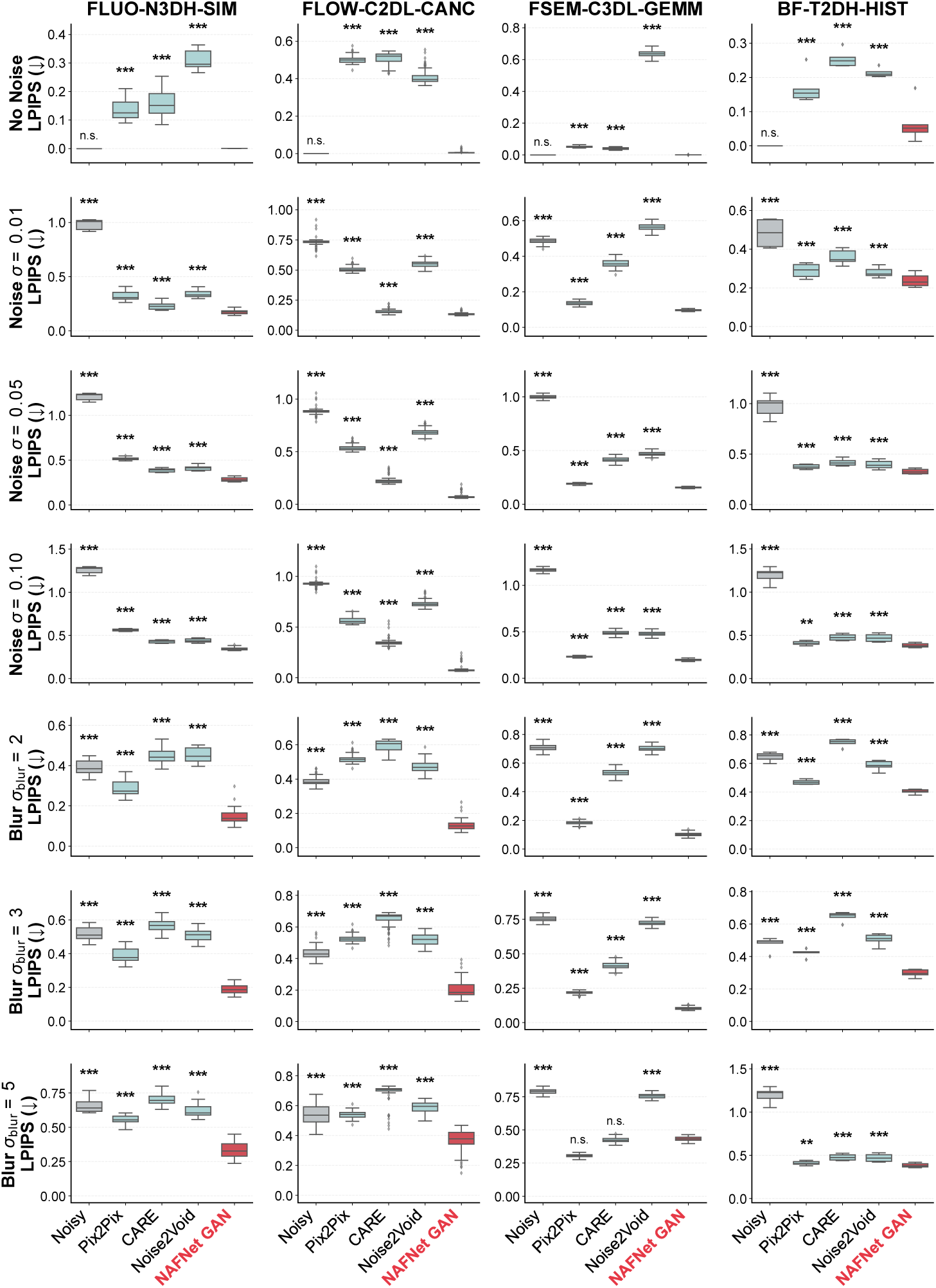
Quantitative stability analysis (Standard Baselines). LPIPS perceptual error trends for NAFNet GAN versus standard baselines (Pix2Pix, CARE, Noise2Void) across four unpaired datasets. Top rows: performance under increasing Gaussian noise (*σ* ∈ {0.00, 0.01, 0.05, 0.10}). Bottom rows: performance under increasing Gaussian blur (*σ*_blur_ ∈ {2, 3, 5}). Statistical significance determined by one-sided Welch’s t-test (* *P <* 0.05; ** *P <* 0.01; *** *P <* 0.001; n.s., not significant).

**Extended Data Fig. 6.**
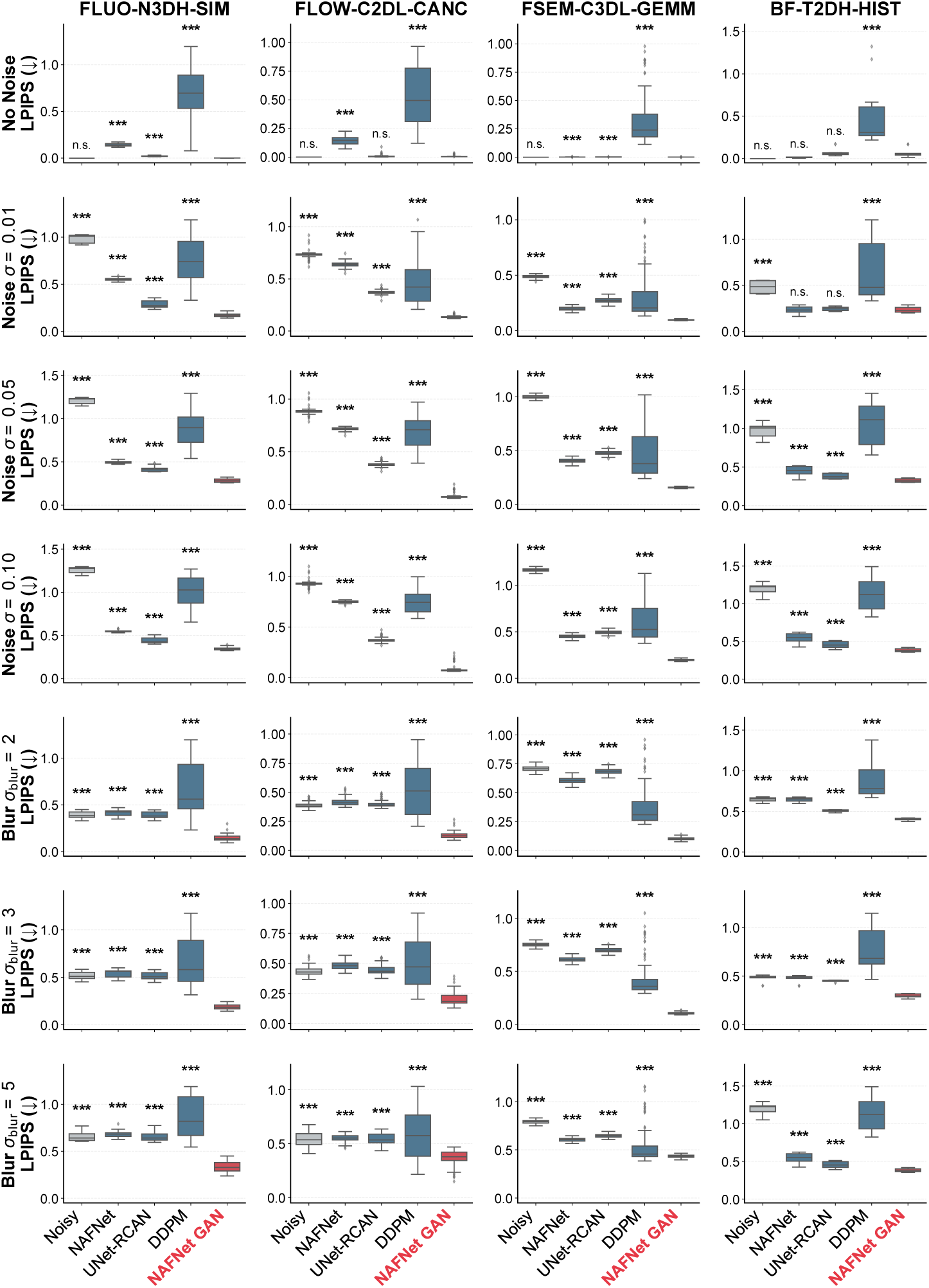
Quantitative stability analysis (Advanced Models). LPIPS perceptual error trends for NAFNet GAN versus advanced architectures (NAFNet, UNet-RCAN, DDPM). Organization matches Extended Data Fig. 5. NAFNet GAN demonstrates significantly lower error variance (*P <* 0.001 in most conditions) across degradation shifts compared to diffusion-based approaches. Statistical significance determined by one-sided Welch’s t-test (* *P <* 0.05; ** *P <* 0.01; *** *P <* 0.001; n.s., not significant).

**Extended Data Fig. 7.**
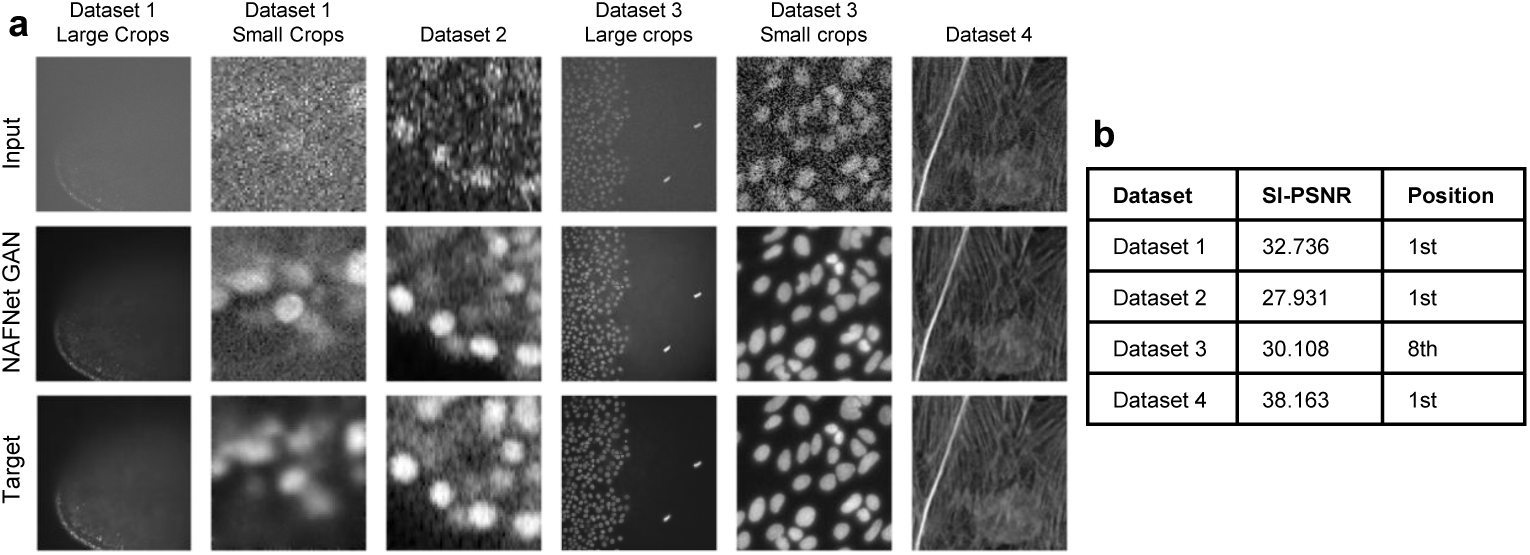
External validation on the AI4Life Microscopy Denoising Challenge. Performance on held-out test sets from the AI4Life 2025 competition. **a**, Representative visual results across four tracks (*Planaria*, Zebrafish, Nuclei, Heterogeneous). **b**, Leaderboard summary. NAFNet GAN ranked **1st** in three categories based on mean Structural Integrity PSNR (SI-PSNR), demonstrating robustness on held-out test sets.

**Extended Data Fig. 8.**
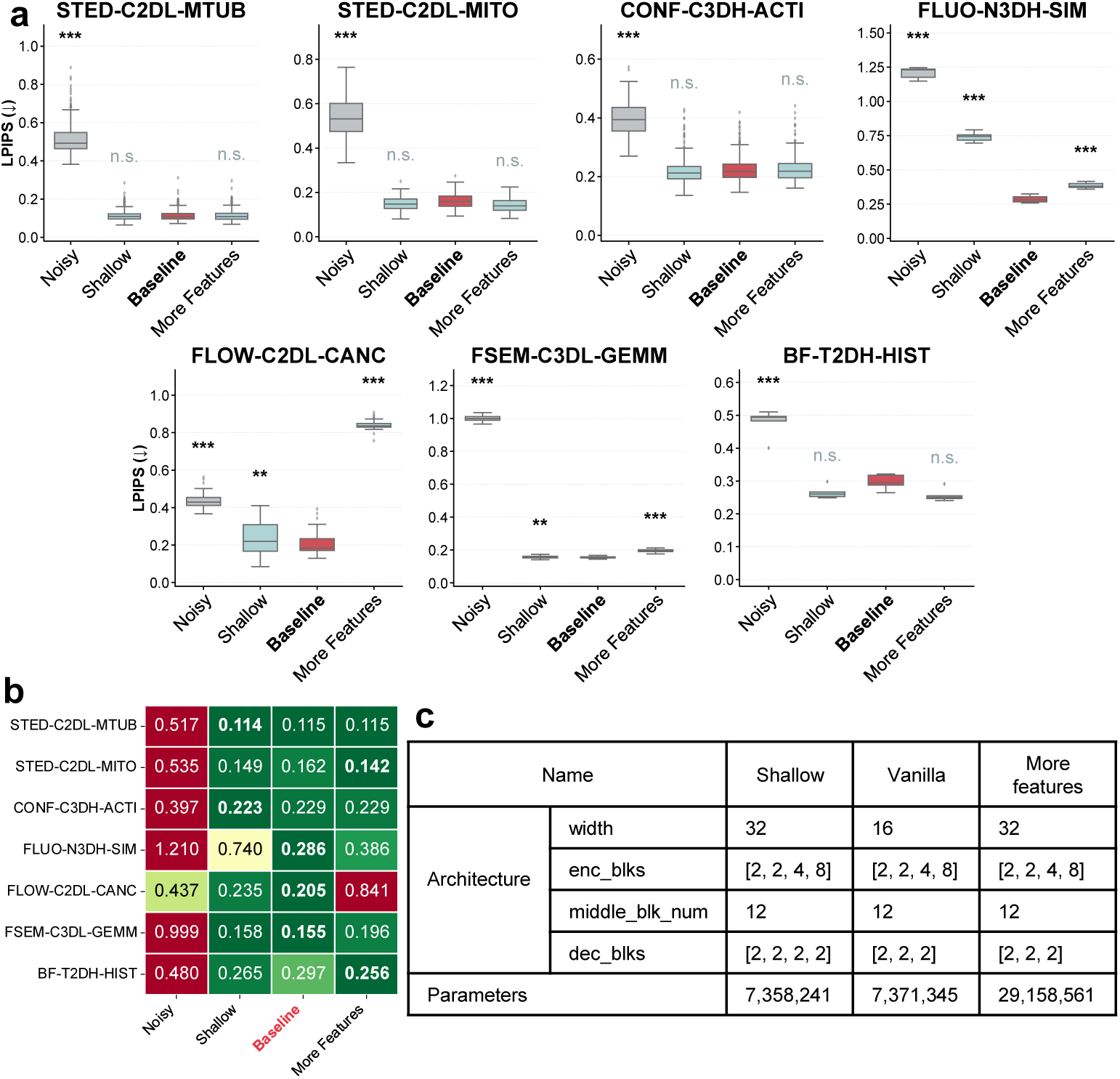
Impact of generator depth on performance. Ablation study comparing the selected Baseline (4-level encoder-decoder) against Shallow (reduced depth) and High-Capacity (increased features) variants. **a**, LPIPS perceptual error across datasets. **b**, Quantitative summary of mean LPIPS scores. **c** Architectural specifications for each variant, including input feature width, NAFNet block distribution (encoding, bottleneck, and decoding), and total parameter counts. The Baseline achieves performance comparable to the High-Capacity model, indicating high efficiency before diminishing returns. Statistical significance determined by one-sided Welch’s t-test (* *P <* 0.05; ** *P <* 0.01; *** *P <* 0.001; n.s., not significant).

**Extended Data Fig. 9.**
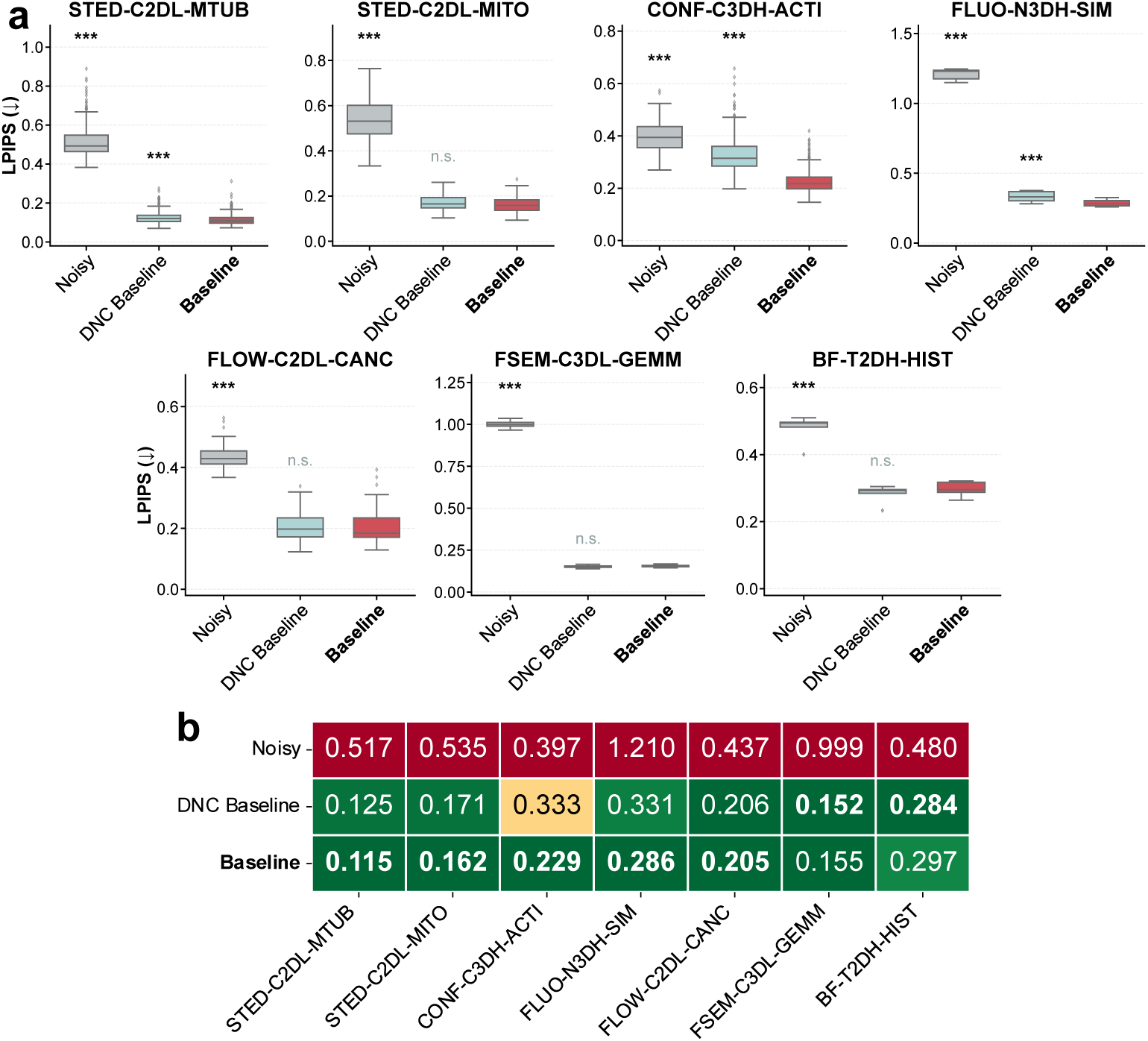
Impact of the optimization objective on perceptual fidelity. Quantitative comparison of two training strategies: (1) Baseline (adversarial loss), and (2) Microscopy Denoising Challenge (DNC) Baseline (challenge-optimized SI-PSNR loss). **a**, LPIPS perceptual error across eight datasets. **b**, Heatmap summary of mean LPIPS scores. The adversarial baseline yields significantly lower perceptual error, confirming the necessity of the GAN objective for texture recovery. Statistical significance determined by one-sided Welch’s t-test (* *P <* 0.05; ** *P <* 0.01; *** *P <* 0.001; n.s., not significant).

